# LIN-35 and the DREAM complex promote temperature stress induced increases in germline apoptosis and cytoplasmic streaming in *C. elegans*

**DOI:** 10.1101/2024.11.13.623436

**Authors:** Frances V. Compere, Kristen A. Quaglia, Margaret N. Crespo Cruz, Hannah N. Lorenzen, Samantha H. Oswald, Katherine Uttal, Lisa N. Petrella

## Abstract

As modest increases in temperature become more common due to global climate change, species are being subjected to moderate temperature stress that can disproportionally affect species fertility. Species that can buffer fluctuations in temperature through tissue or cellular responses in the germline will therefore be more likely to survive moderate temperature stress. Currently, what mechanisms are used in the germline to facilitate maintenance of fertility under moderate temperature stress are unknown. To address this, we investigated how germline apoptosis is modulated in *Caenorhabditis elegans* nematodes in response to moderate temperature stress. We found that wildtype animals increase their germline apoptosis levels from the physiological baseline in response to the moderate temperature stress. This induction of germline apoptosis was dependent on conserved members of the DREAM complex, including LIN-35, the *C. elegans* homolog of the retinoblastoma tumor and members of the MuvB core, LIN-54 and LIN-37. We also found that LIN-35, LIN-54, and LIN-37 were necessary for DNA damage induced apoptosis. Additionally, induction of germline apoptosis during moderate temperature stress was dependent on repression CED-9 function, the *C. elegans* Bcl2 ortholog. Finally, we found that changes in cytoplasmic streaming likely lead to changes to oocyte provisioning in wildtype animals but not mutants. Together, these data suggest an expanded role for LIN-35, CED-9, and the DREAM complex in maintaining fertility by activating apoptosis during moderate temperature stress.

## INTRODUCTION

For a species to maintain population size, organisms must respond to environmental stressors to maintain fertility. Given the unique contributions of oocytes to both the genome and cytoplasm of embryos, the oogenic germline plays a major role in ensuring fertility and progeny fitness under stressful conditions. Apoptosis has been shown to be a mechanism used in the oogenic germline to respond to a number of different stressors in *C. elegans* including DNA damage, osmotic stress, oxidative stress, starvation, ethanol stress, and aging (Andux and Ellis 2008, Fausett *et al*. 2021; Gartner *et al*. 2000; Salinas *et al*. 2006). However, how changes in the level of apoptosis contribute to maintaining fertility during the abiotic stress of moderately increased temperature remains unexplored. Moderate increases in temperature are known to negatively affect fertility in a wide range of organisms including mammals, plants, insects, and nematodes (Zinn *et al*. 2010; Takahashi 2011; Petrella 2014; Gandara and Drummond-Barbosa 2023; Tushabe *et al*. 2023). Given the projected continuing increases in temperatures with global climate change, it is imperative to understand which mechanisms contribute to the maintenance of fertility during moderate temperature stress.

In *C. elegans* hermaphrodites, the germline in adults is oogenic and the sole site of post-developmental apoptosis (Gartner *et al*. 2008). Hermaphrodites contain two gonad arms with a shared uterus. Each gonad is a u-shaped tube that contains germ cells surrounded by a thin layer of supporting somatic cells (Hubbard and Greenstein 2005). In the section of the germline distal to the uterus, the germline nuclei are only partially surrounded by a cell membrane and thus are in a syncytium with a shared core of cytoplasm called the rachis. By convention these nuclei are termed germ cells despite being only partially cellularized (Raiders *et al*. 2018). Germ cells initially undergo mitosis and then transition into meiosis as they move down the gonad. During non-stressful conditions, approximately 50% of germ cells undergo what is termed physiological apoptosis as they approach the bend of the gonad (Gumienny *et al*. 1999). These dead germ cells are then engulfed by the surrounding somatic sheath cells and most of the associated cytoplasm and organelles are directed to the remaining living nuclei through cytoplasmic streaming (Ellis *et al.,* 2001; Wolke *et al*. 2007). Cytoplasmic streaming leads the nuclei destined to become mature oocytes obtaining a large store of cytoplasm/organelles as they cellularize (Wolke *et al*. 2007).

Stress conditions that lead to nuclear damage trigger the removal of damaged nuclei through apoptosis, which leads to a higher level of apoptosis than occurs during non-stress conditions (Schumacher *et al*. 2001; Bhalla and Dernburg 2005). Thus, apoptosis acts as a cellular response in the oogenic germline to environmental stress resulting in two related outcomes that may help to maintain fertility (Gartner 2008; Cao and Pocock 2022). The first outcome is the removal of damaged nuclei to limit the inheritance of damaged genomes, and the second outcome is a potential increase in the amount of cytoplasm the germ line contributes to oocytes that may impact early embryonic health. Like other stressors, moderate temperature stress of 26-27°C in *C. elegans* also has been shown to increase the level of germline apoptosis (Poullet *et al*. 2015). At this temperature range *C. elegans* goes from having few progeny (26°C) to being basically sterile (27°C) (Petrella 2014). While the molecular mechanisms leading to increased levels of apoptosis are well understood for other stressors, the molecular mechanisms leading to increased apoptosis under moderate temperature stress are unknown.

Stress induced germline apoptosis in *C. elegans* uses the same conserved signaling pathway as is found across eukaryotes (Ellis and Horvitz 1986; Gartner *et al*. 2008). CED-9, the *C. elegans* Bcl-2 homolog, functions to repress activation of the core apoptotic caspase machinery by sequestering CED-4, the *C. elegans* Apaf homolog (Hengartner *et al*. 1992; Spector *et al*. 1997; Chen *et al*. 2000). When the CED-9 protein function is inhibited, CED-4 is released and goes on to activate CED-3, the terminal caspase that activates apoptosis (Chinnaiyan *et al*. 1997; Spector *et al*. 1997; Chen *et al*. 2000). LIN-35, the sole *C. elegans* homolog of the mammalian pocket proteins, has been shown to be necessary for the increase in germline apoptosis under starvation and DNA damage conditions (Schertel and Conradt 2007; Láscarez-Lagunas *et al*. 2014). LIN-35 has been shown to modulate apoptosis levels through the repression of *ced-9* mRNA expression (Schertel and Conradt 2007; Láscarez-Lagunas *et al*. 2014). However, LIN-35 does not have DNA binding activity and its repressive activity is generally mediated through its interactions within protein complexes. The most well studied LIN-35 interaction is with the conserved DREAM complex (Dp, Retinoblastoma (Rb)-like, E2F, MuvB) (Harrison *et al*. 2006; Goetsch *et al*. 2017). The DREAM complex is made up of a E2F/DP dimer linked through a pocket protein, such as LIN-35, with the Muv B core (Harrison *et al*. 2006; Goetsch *et al*. 2017). The Muv B core in *C. elegans* is made up of five proteins, including LIN-54 and LIN-37. No studies to date have looked at a role for the Muv B core of the DREAM complex in regulation of *C. elegans* germline apoptosis. However, the conserved members of the DREAM complex, including LIN-35, have been shown in embryos to bind to the *ced-9* operon (Goetsch *et al*. 2017). Therefore, members of the Muv B core of the DREAM complex are potential cofactors for LIN-35 in induction of increased germline apoptosis during stress, including moderate temperature stress.

Here we investigate the mechanisms leading to increased apoptosis during moderate temperature stress and explore the effects of moderate temperature stress on cytoplasmic streaming, oocyte size and ovulation rate. We find that the increase in apoptosis during moderate temperature stress depends on both LIN-35 and the Muv B core in addition to repression of CED-9 function. During moderate temperature stress there is also an increase in cytoplasmic streaming in wild type with a concomitant increase in oocyte size. Overall, these findings further underscore LIN-35 and the Muv B core of the DREAM complex as general regulators of stress-induced germline apoptosis. In turn, increased apoptosis during moderate temperature stress provides mechanism for increased oocyte size that may be protective for embryonic development.

## METHODS

### Strains and nematode culture

*C. elegans* were cultured under standard conditions (Brenner 1974) on NGM plates seeded with *E. coli* strain AMA1004 at 20°C unless otherwise noted. N2 was used as the wild type control unless otherwise noted. Strains used in this study were N2, MT8841 *lin-54(n2231) IV,* MT10430 *lin-35(n745) I,* MT5470 *lin-37(n758) III*, MD701 *bcls39(lim-7p::ced-1::GFP) V,* LNP0089 *lin-35(n745); bcls39(lim-7p::ced-1::GFP) V,* LNP0091 *lin-54(n2231) IV; bcls39(lim-7p::ced-1::GFP) V,* LNP0092 *lin-37(n768) III; bcls39(lim-7p::ced-1::GFP) V*, LNP0266 *ced-9(n1950) III; bcls39(lim-7p::ced-1::GFP) V,* LNP0268 *ced-9(n1950) III; lin-54(n2331) IV*; *bcls39(lim-7p::ced-1::GFP) V*. All LNP strains were made for this study. The N2 strain was from the laboratory of Susan Strome. All other strains were provided by the CGC, which is funded by NIH Office of Research Infrastructure Programs (P40 OD010440).

### CED-1::GFP apoptosis assay

#### Treatments

*Control for temperature treatments:* Worms were grown to the L4 stage at 20°C, then isolated and maintained at 20°C for 24 hours prior to imaging. Three or four biological replicates were performed, with 5-10 worms analyzed per replicate for a total of n = 50-54 germline arms per genotype.

*Upshift to 26°C at L1 stage:* Worms were grown to the L4 stage at 20°C, then 10- 20 P0 L4 worms were isolated and maintained at 20°C for 24 hrs until they all reached the adult stage. All P0 worms were moved to new plates and allowed to lay embryos for 3 hours before being removed. F1 embryos were allowed to hatch and grow for 24hrs at 20°C, then upshifted to 26°C until the L4 stage. F1 L4 worms were then isolated and maintained at 26°C for 24 hours prior to imaging. Two biological replicates were performed, with 8-10 worms analyzed per replicate for a total of n = 34-38 germline arms per genotype.

*Upshift to* 26°C *at L4 stage:* Worms were grown to the L4 stage at 20°C, then isolated and upshifted to 26°C for 24 hours prior to imaging. Three biological replicates were performed, with 5-10 worms analyzed per replicate for a total of n = 40-58 germline arms per genotype.

*Control for UV treatment:* Worms were grown to the L4 stage at 20°C, then isolated and maintained at 20°C for 48 hours prior to imaging. Eight biological replicates were performed, with 5-10 worms analyzed per replicate for a total of n = 154-160 germline arms per genotype.

*UV treatment*: Worms were grown to the L4 stage at 20°C, then isolated and maintained at 20°C for 24 hours prior to UV treatment. Worms were subjected to 400 J/m^2^ of UV in a CL-1000 Ultraviolet Crosslinker and then allowed to recover for 24 hours at 20°C prior to imaging. Eight biological replicates were performed, with 5-10 worms analyzed per replicate for a total of n = 148-160 germline arms per genotype.

#### Apoptosis scoring and statistical analysis

Worms were mounted on a 2% agarose pad in 10uM levamisole in 1X egg buffer. Each germline arm was scored for GFP positive apoptotic cells using the *ced-1::GFP* transgene on a Nikon Eclipse TE2000-S inverted microscope equipped with a Plan Apo 60X/1.25 numerical aperture oil objective. Statistical analysis was done using a two-way ANOVA with Tukey’s correction using Prism 10.0.3 (GraphPad Boston, MA).

### Oocyte Size and Number Analysis

Worms were grown to the L4 stage at 20°C, then isolated and maintained at 20°C or upshifted to 26°C for 24 hours prior to imaging. For imaging, worms were mounted on slides with 2% agarose pads in 10uM levamisole in 1X M9, without bacteria. Oocytes from one gonad arm were imaged from each worm using a Nikon Eclipse TE2000-S inverted microscope equipped with a Plan Apo 60x/1.25 numerical aperture oil objective. Images were captured using a Q imaging Exi Blue camera (Teledyne Photometrics, Tucson AZ, USA) using Nomarski optics and Q Capture Pro 7 software (Teledyne Photometrics, Tucson AZ, USA). The area of each fully cellularized oocyte within one germline arm was measured in FIJI using the freehand tool with a scale of 4.64in by 3.47in (Schindelin *et al*. 2012). The area was measured by tracing around the membrane of each of the fully cellularized oocytes. Oocytes were considered fully cellularized if they had a cell membrane fully surrounding the nucleus. Three to five biological replicates were performed, with 3-6 worms analyzed per replicate for a total of n= 17-27 germline arms per genotype per temperature. Statistical analysis was done using a Two-Way ANOVA with Tukey’s multiple comparisons using Prism 10.0.3 (GraphPad Boston, MA).

### Ovulation Rate Analysis

Ovulation rate analysis was performed as in Rios et al. (2017). Worms were grown to the L4 stage at 20°C, then isolated and maintained at 20°C or upshifted to 26°C for 24 hours prior to imaging. On the day of the ovulation assay, adult worms were cloned to a plate with only a thin lawn of *E. coli* and the embryos inside the uterus were immediately counted. Worms were then allowed to lay embryos for 3 hours at the appropriate temperature. At the end of the three hours the embryos inside the uterus were counted, then the worm was removed from the plate, and the embryos laid on the plate were counted. All steps were visualized on a Nikon SMZ1500 stereomicroscope. We used the following formula to calculate the ovulation rate per gonad arm per hour using the formula ((Final # embryos in uterus - Initial # embryos in uterus) + Number of embryos on the plate)/(2 gonads X 3 hours). Eight to 10 biological replicates were performed, with 1-3 worms analyzed per replicate for a total of n = 16-26 worms per genotype. Statistical analysis was done using a two-way ANOVA using with Tukey’s correction Prism 10.0.3 (GraphPad Boston, MA).

### Cytoplasmic Streaming Analysis

Cytoplasmic streaming analysis was performed as in Wolke et al (2007). Worms were grown to the L4 stage at 20°C, then isolated and maintained at 20°C or upshifted to 26°C for 24 hours prior to imaging. Immediately before imaging, all worms were anesthetized in a 1μM levamisole in 1X M9 buffer solution for 5 minutes in a glass well and then placed on a 4% agarose pad while maintained in the levamisole. Worms were imaged every 15 seconds over 20 min using Nomarski optics on a Nikon Eclipse TE2000-S inverted microscope equipped with a Plan Apo 60X/1.25 numerical aperture oil objective. Images were captured using a Q imaging Exi Blue camera (Teledyne Photometrics, Tucson AZ, USA) and Q Capture Pro 7 software (Teledyne Photometrics, Tucson AZ, USA). To measure the speed of cytoplasmic streaming, particles within the rachis were chosen that were approaching or traveling around the germline bend and were visible within the focal plane for a minimum of 2 min. Each particle was tracked over 2 minutes using the “Manual Tracking” plugin in Fiji (Schindelin *et al*. 2012). For each genotype and temperature five particles in each of six independent germlines were analyzed. Statistical analysis was done using Two-way ANOVA with Tukey’s correction using Prism 10.0.3 (GraphPad Boston, MA).

### Data Availability Statement

Strains and plasmids are available upon request. The authors affirm that all data necessary for confirming the conclusions of the article are present within the article, figures, and tables.

## RESULTS

### lin-35 and DREAM complex mutants do not fully induce germline apoptosis in response to moderate temperature stress

Given the known roles of LIN-35 in promoting germline apoptosis under other conditions (Schertel and Conradt 2007; Láscarez-Lagunas *et al*. 2014), we investigated whether LIN-35 promotes increased germline apoptosis levels during moderate temperature stress at 26°C. As LIN-35 is known to interact with the DREAM complex and has been shown to co-bind with the DREAM complex at the *ced-9* operon (Goetsch *et al*. 2017), we also investigated whether members of the Muv B core of the DREAM complex, LIN-54 and LIN-37, could also promote germline apoptosis (Figure 1A). To determine the level of apoptosis, we used strains that carried the CED-1::GFP transgene, which allows for visualization of germline apoptotic cells (Zhou *et al*. 2001; Schumacher *et al*. 2005). We counted apoptotic cells in wild type and the three mutant strains *lin-35(n745), lin-54(n2231), lin-37(n758)* under three temperature conditions: continual exposure to 20°C (non-stress condition), up-shifting to 26°C at the L1 stage (developmental temperature stress), and up-shifting to 26°C at the L4 stage (post-germline development temperature stress). We assessed the number of apoptotic cells in worms 24 hrs post L4 stage. Wildtype, but not mutant, hermaphrodites showed a significant increase in apoptosis when worms were upshifted at the L1 stage compared to the same genotype maintained at 20°C (Figure 1B). On the other hand, all genotypes showed a significant increase in apoptosis when hermaphrodites were upshifted at the L4 stage compared to the same genotype maintained at 20°C (Figure 1B). However, all three mutants showed a significantly lower level of apoptosis than wild type for both 26°C stress conditions (Figure 1B). These data suggest a role for LIN-35 and the DREAM complex Muv B core in the induction of germline apoptosis in response to moderate temperature stress.

**Figure 1:**
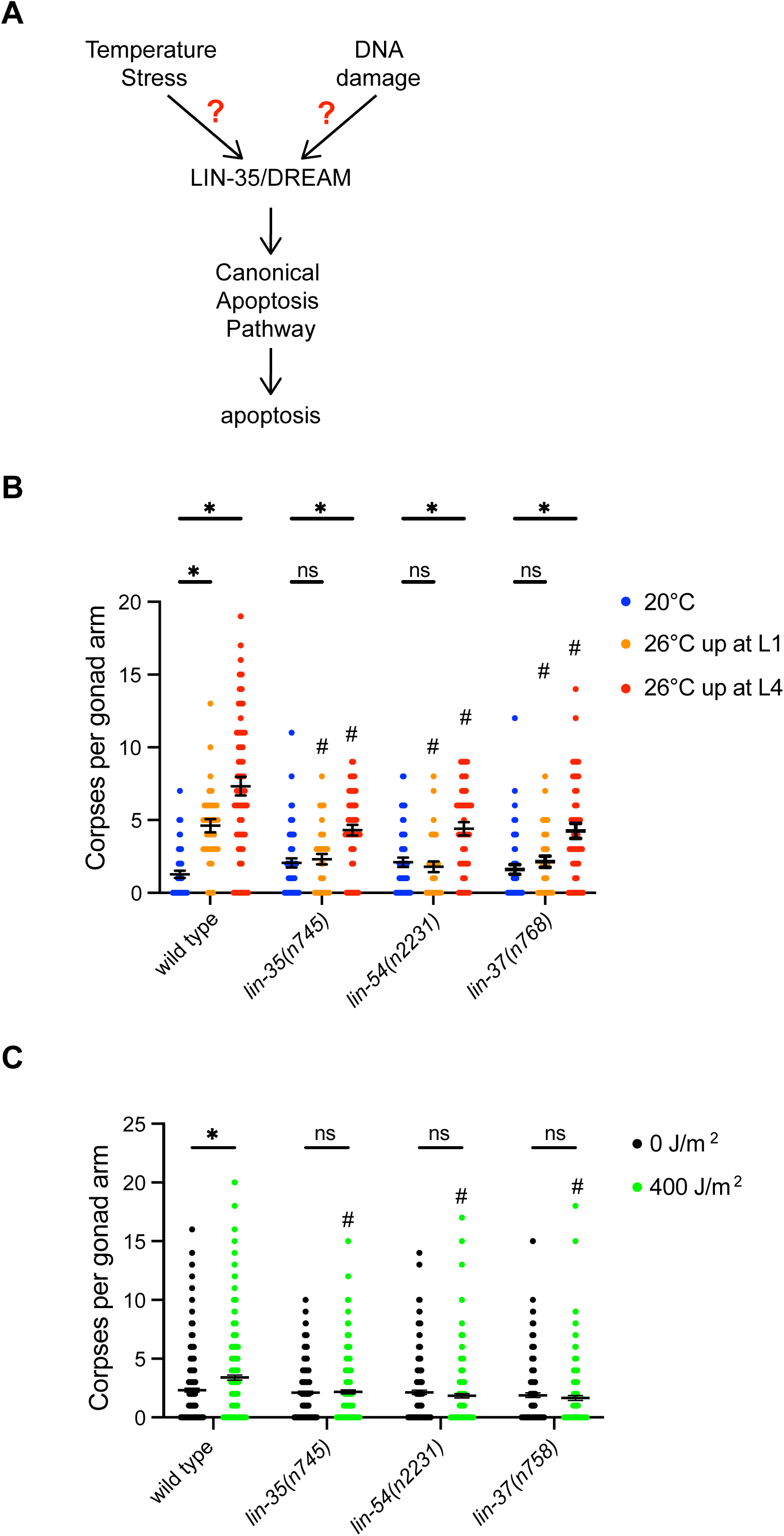
LIN-35 and the DREAM complex were necessary for apoptosis induction during temperature stress and DNA damage. (A) Model where moderate temperature stress or DNA damage could work through LIN-35 and the DREAM complex to activate germline apoptosis through the canonical apoptosis pathway. (B) Germ cell corpses counted per gonad arm using *ced-1::gfp* in hermaphrodites 24 hours post-L4 stage in the indicated genetic backgrounds. Hermaphrodites were either maintained continually at 20°C (blue), upshifted to 26°C at the L1 stage (orange), or upshifted to 26°C at the L4 stage (red) with each dot representing an individual gonad, n = 34-58 gonads. * significantly different within the genotype compared to 20°C, # significantly different than wild type at the same temperature. *P-*value ≤ 0.05 using two-way ANOVA with Tukey correction. Error bars indicate ± SEM. (C) Germ cell corpses counted per gonad arm using *ced-1::gfp* in hermaphrodites 48 hours post-L4 stage in the indicated genetic backgrounds. Hermaphrodites 24 hours post-L4 were either treated with 0 J/m^2^ (black) or 400 J/m^2^ (green) of UV then allowed to recover for 24 hours, n = 168-298 gonads. * significantly different within the genotype compared to 0 J/m^2^, # significantly different than wild type with the same UV treatment. *P-*value ≤ 0.05 using two-way ANOVA with Tukey correction. Error bars indicate ± SEM.

### lin-35 and DREAM complex mutants do not induce germline apoptosis in response to UV induced DNA damage

Previous studies have shown that LIN-35 is important for DNA damage induced germline apoptosis (Schertel and Conradt 2007); however, it was unknown whether this was in conjunction with its role in the DREAM complex. Therefore, we investigated if the DREAM complex Muv B core mutants, *lin-54(n2231)* and *lin-37(n758)*, had reduced induction of germline apoptosis in response to UV induced DNA damage similar to *lin-35* mutants. We found that, as expected, wild type animals demonstrated an increase in the number of apoptotic cells when subjected to 400 J/m^2^ of UV radiation compared to wildtype animals that were not subjected to UV (Figure 1C). None of the mutants showed an increase of in the number of apoptotic cells when subjected to 400 J/m^2^ of UV radiation compared to the same genotype that were not subjected to UV (Figure 1C). Like with temperature stressed worms, all three mutants also showed fewer apoptotic cells during UV treatment than wild type (Figure 1C). These results strongly suggest that the DREAM complex may function with LIN-35 in the germline to help promote increased germline apoptosis under a variety of stressors, but these proteins do not appear to be necessary to regulate physiological apoptosis.

### Constitutively active CED-9 mutants do not induce apoptosis to wild type levels in response to moderate temperature stress

Previous research has shown that regulation of *ced-9* expression by *lin-35* during DNA damage and starvation apoptosis (Schertel and Conradt 2007; Láscarez-Lagunas *et al*. 2014). This suggests that *ced-9* mRNA levels, and also likely CED-9 protein levels, could play a role in induction of germline apoptosis during multiple stress conditions. To further investigate the potential role of CED-9 regulation in stress induce germline apoptosis we used the *ced-9(n1950)* mutation, which constitutively sequesters CED-4 leading to a block in activation of apoptosis (Figure 2A) (Hengartner and Horvitz 1994). Above we found that upshifting hermaphrodites to 26°C at the L4 stage had the strongest effect on the activation of germline apoptosis (Figure 1B), therefore we used this temperature treatment for all of our subsequent experiments. We found that *ced-9(n1950)* mutants showed a smaller increase in apoptosis compared to wild type animals, similar to that seen in *lin-35* and DREAM complex mutants with the same temperature treatment (Figure 1B and 2B). We next measured the level of apoptosis in *ced-9(n1950); lin-54(n2231)* double mutants using the same temperature treatment. We found that the double mutant had no induction of apoptosis during moderate temperature stress (Figure 2B). Consistent with previously published data (Gumienny *et al*. 1999), the level of apoptosis did not go to zero in either mutant containing *ced-9(n1950)* under any conditions. Thus, similar to other stresses that have been tested (Salinas *et al*. 2006; Láscarez-Lagunas *et al,* 2014), repression of CED-9 function is important for the increase in apoptosis during moderate temperature stress, but not for physiological apoptosis.

**Figure 2:**
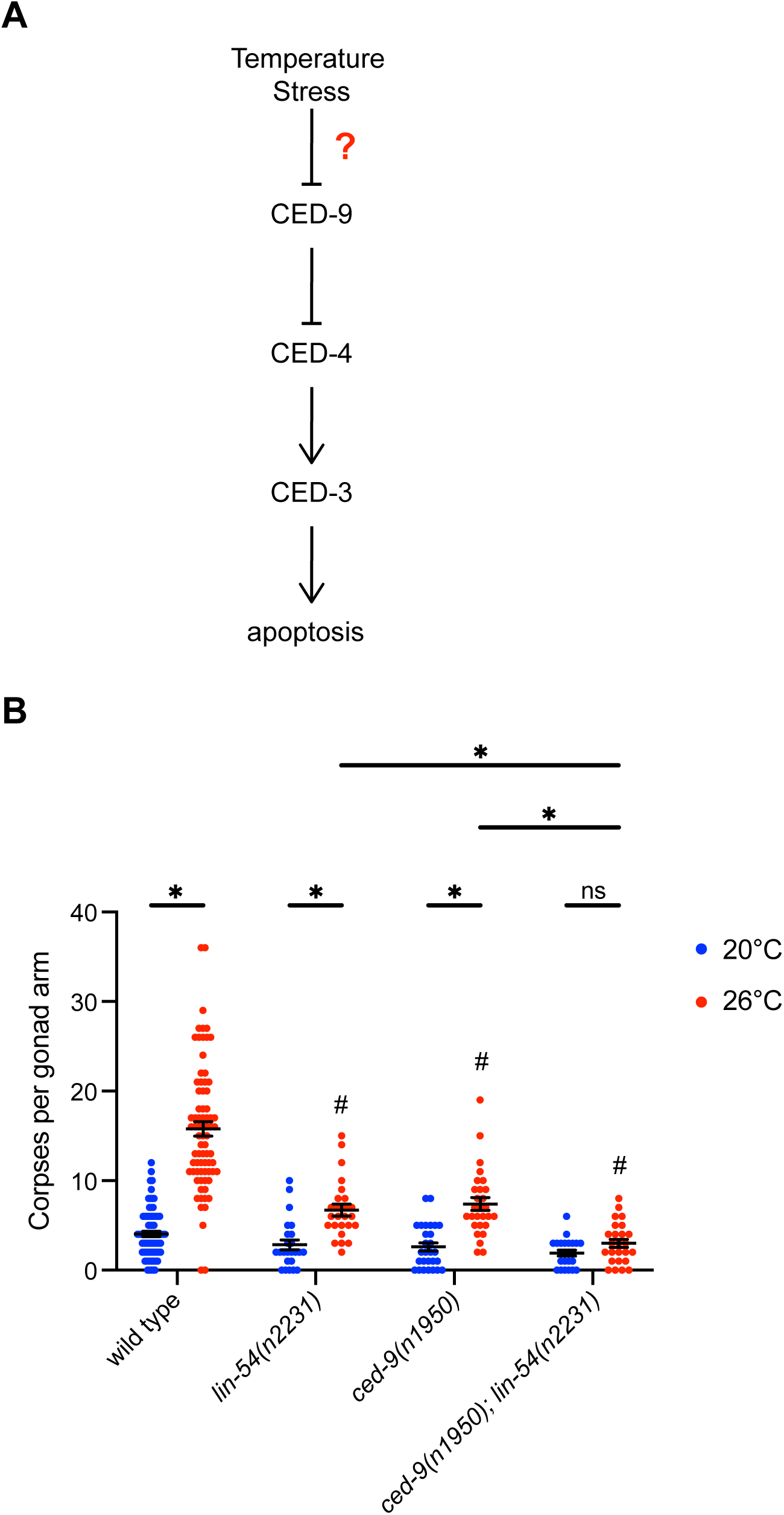
Repression of CED-9 function is necessary for apoptosis induction during temperature stress. (A) Model of the canonical apoptosis pathway in *C. elegans.* CED-9 functions to sequester CED-4 away from CED-3. When apoptosis is induced, potentially through moderate temperature stress, CED-9 function is suppressed, leading to CED-4 activation of CED-3 leading to apoptosis. (B) Germ cell corpses counted per gonad arm using *ced-1::gfp* in hermaphrodites 24 hours post-L4 stage in the indicated genetic backgrounds. Hermaphrodites were either maintained continually at 20°C (blue) or upshifted to 26°C at the L4 stage (red) with each dot representing an individual gonad, n = 24-78 gonads. * significantly different within the genotype compared to 20°C or between a single mutant and double mutant within the same temperature, # significantly different than wild type at the same temperature. *P-* value ≤ 0.05 using two-way ANOVA with Tukey correction. Error bars indicate ± SEM.

### lin-35 and DREAM complex mutants did not show increased cytoplasmic streaming in response to moderate temperature stress

One of the functions of germline apoptosis is the contribution of cytoplasmic components from the dying nuclei to developing oocytes (Gartner *et al*. 2008). In *C. elegans,* cytoplasm that is expelled from dying nuclei into the central core of shared cytoplasm (rachis) moves around the bend of the germline into cellularizing oocytes in a process called cytoplasmic streaming (Figure 3A; Wolke *et al*. 2007). In addition, while apoptosis is not required for cytoplasmic streaming to occur, the rate of cytoplasmic streaming has been shown to depend on apoptosis (Wolke *et al*. 2007). Since temperature generally increases the rate of many cellular and physiological processes, we investigated changes in cytoplasmic streaming in wild type, *lin-35(n745),* and *lin-54(n2231)* mutants at different temperatures. We found that the rate of cytoplasmic streaming was significantly higher in wild-type worms upshifted to 26°C compared to wild-type worms maintained at 20°C (Figure 3B). In contrast, we found no change in the rate of cytoplasmic streaming in *lin-35(n745)* or *lin-54(n2231)* mutants upshifted to 26°C compared to worms maintained at 20°C (Figure 3B). This failure to increase cytoplasmic streaming in the mutants when upshifted to 26°C corresponded with their failure to increase apoptosis to the same level as wild type under the same temperature conditions. We also found that *lin-54(n2231)* mutants had a significantly higher rate of cytoplasmic streaming at 20°C than that of wild type at 20°C (Figure 3B). On the other hand, both *lin-35(n745)* and *lin-54(n2231)* mutants had significantly slower rates of cytoplasmic streaming when upshifted to 26°C than that of wild type upshifted to 26°C (Figure 3B). These data support a model where increased apoptosis under temperature stress contributes to increased cytoplasmic streaming. Increased cytoplasmic streaming during temperature stress could in turn lead to oocytes similar in sized to those in unstressed animals but accompanied with a faster ovulation rate or the production of larger oocytes than in unstressed animals.

**Figure 3:**
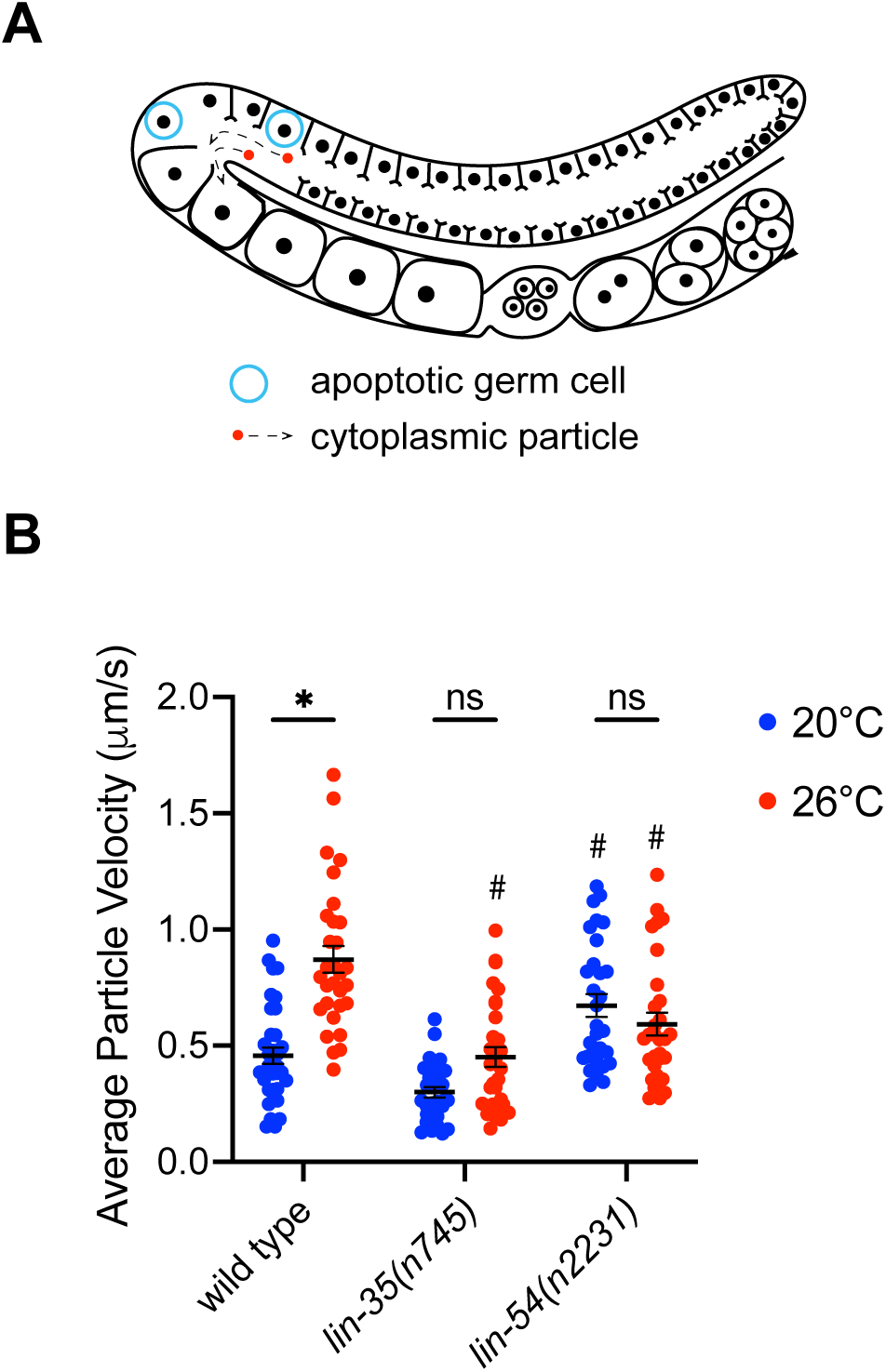
LIN-35 and the DREAM complex were necessary to increased cytoplasm streaming at during temperature stress. (A) Model of one arm of a oogenic germline. Apoptotic germ cells being engulfed are represented by blue circles. Cytoplasmic streaming represented by the trajectory of the red dots. (B) The rate of cytoplasmic particle movement was measured under DIC microscopy in wild type, *lin-35(n745),* and *lin-54(n2231)* mutants. Hermaphrodites were either maintained continually at 20°C (blue) or upshifted to 26°C at the L4 stage (red) with each dot representing an individual particle tracked, n = 5 particles per 6 gonads for a total 30 particles per genotype tracked. * significantly different within the genotype compared to 20°C, # significantly different than wild type at the same temperature. *P-*value ≤ 0.05 using two-way ANOVA with Tukey correction. Error bars indicate ± SEM.

### Temperature does not affect ovulation rate in wild type or mutants

We investigated if ovulation rate increased when worms are exposed to post-developmental temperature stress. We found that neither wild type nor mutant adult worms demonstrated an increase in ovulation rate when upshifted to 26°C at the L4 stage compared to the same strain at 20°C (Figure 4). However, both *lin-35(n745)* and *lin-54(n2231)* mutants had significantly slower ovulation rates than wild type at the same temperature (Figure 4). These data suggest that neither temperature nor apoptosis level directly affect the rate of ovulation. However, the pleotropic defects in the germline experienced in *lin-35(n745)* and *lin-54(n2231)* mutants (Goetsch *et al.,* 2019; Mikeworth *et al*. 2023) seem to affect overall ovulation rate.

**Figure 4:**
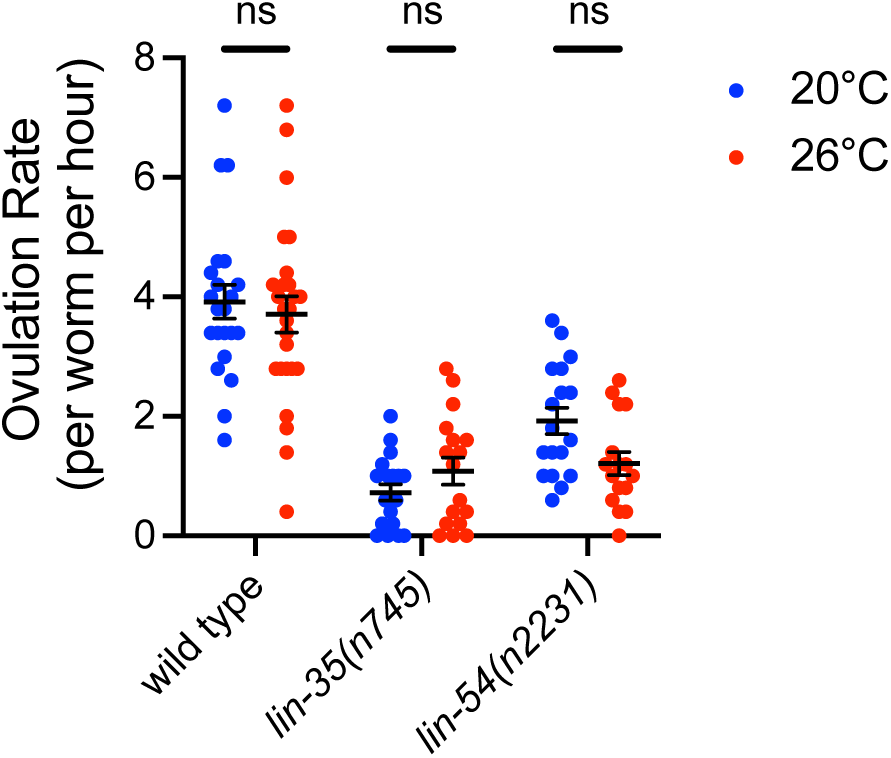
Ovulation rate does not change in wild type, *lin-35* or DREAM complex mutants during temperature stress. Ovulation rates measured in wild type, *lin-35(n745),* and *lin-54(n2231)* mutants. Hermaphrodites were either maintained continually at 20°C (blue) or upshifted to 26°C at the L4 stage (red) with each dot representing the ovulation rate within an individual worm, n = 16-26 worms per genotype. ns = not significantly different within the genotype compared to 20°C. *P-* value ≤ 0.05 using two-way ANOVA with Tukey correction. Error bars indicate ± SEM.

### Oocyte size increases with moderate temperature stress

We next investigated if the number or size of oocytes was affected when worms were exposed to post-developmental temperature stress. We found that neither wild type nor mutant adult worms demonstrated a change in the number of oocytes present when upshifted to 26°C at the L4 stage compared to the same strain at 20°C (Figure 5A). However, both *lin-35(n745)* and *lin-54(n2231)* did show a non-significant trend towards fewer oocytes when upshifted to 26°C at the L4 stage compared to the same strain at 20°C (Figure 5A). In addition, both *lin-35(n745)* and *lin-54(n2231)* mutants had significantly lower numbers of oocytes than wild type at the same temperature (Figure 5A). We also found that both wild type and mutant worms had on average larger oocytes when upshifted to 26°C at the L4 stage compared to the same strain at 20°C (Figure 5B). However, *lin-35(n745)* mutants had significantly smaller oocytes than wild type at the same temperature (Figure 5B).

**Figure 5:**
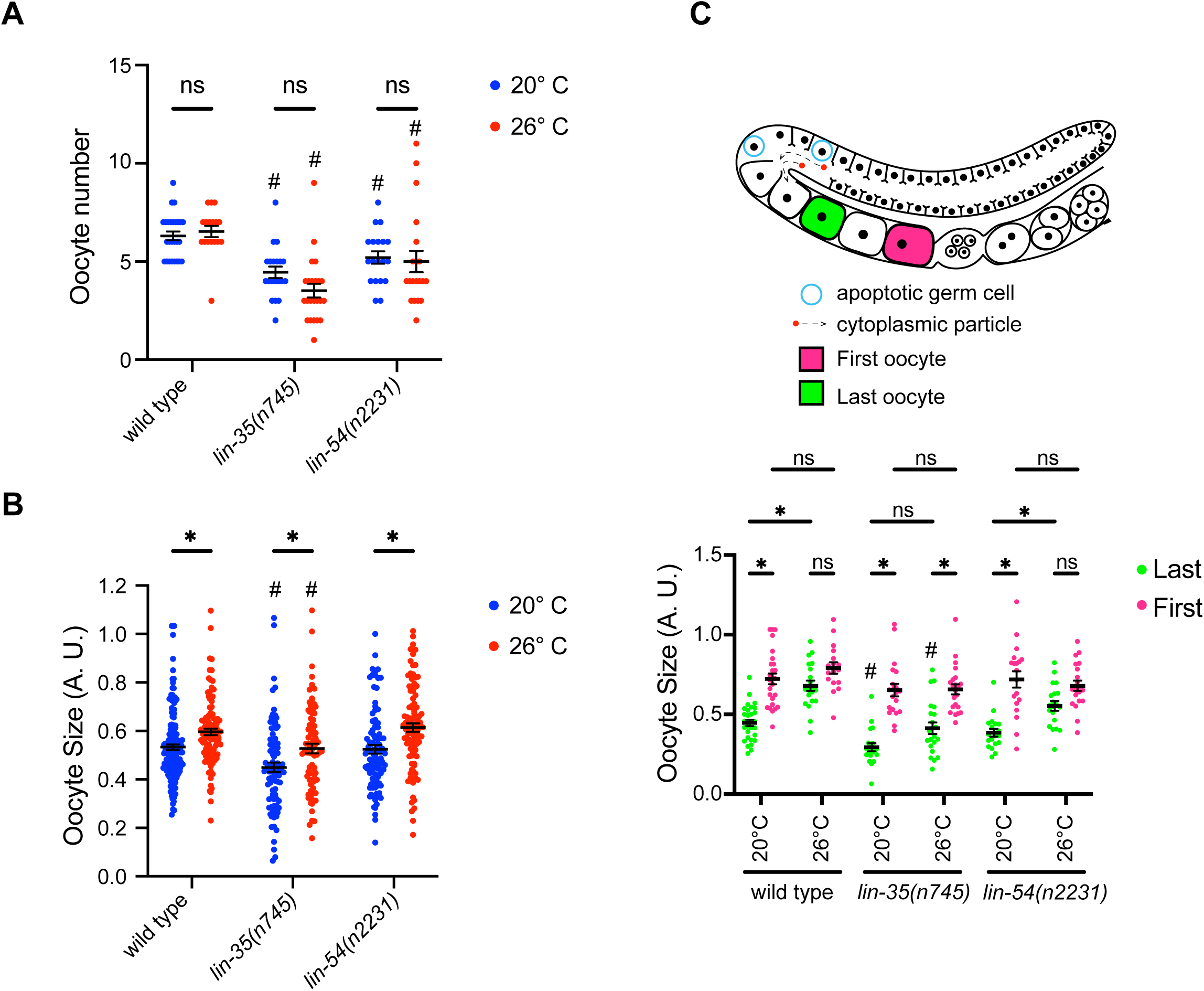
Oocyte size but not number change in wild type, *lin-35* and DREAM complex mutant during temperature stress. (A) Oocyte number was scored in wild type, *lin-35(n745),* and *lin-54(n2231)* mutants. Hermaphrodites were either maintained continually at 20°C (blue) or upshifted to 26°C at the L4 stage (red) with each dot representing the oocyte number within a gonad arm, n = 15-26 worms per genotype per temperature. (B) Individual oocyte sizes were measured in wild type, *lin-35(n745),* and *lin-54(n2231)* mutants. Hermaphrodites were either maintained continually at 20°C (blue) or upshifted to 26°C at the L4 stage (red) with each dot representing an individual oocyte size, n = 81-165, across 15-26 worms per genotype per temperature. *significantly different within the genotype compared to 20°C, # significantly different than wild type at the same temperature. *P-*value ≤ 0.05 using two-way ANOVA with Tukey correction. Error bars indicate ± SEM. (C) Model of the germline with the last (green) and first (pink) oocytes labeled. The measurements of the last (green) and first (pink) oocyte sizes were pulled out of the data from (B) for wild type, *lin-35(n745),* and *lin-54(n2231)* mutants at 20°C or upshifted to 26°C at the L4 stage (red) with each dot representing an individual oocyte size, n =17-30, across 15-26 worms per genotype per temperature. * significantly different *P-*value ≤ 0.05 using two-way ANOVA with Tukey correction. Error bars indicate ± SEM.

Oocyte growth occurs through two mechanisms. The first is that oocytes gain a large portion of their size as they cellularize through cytoplasmic streaming (Wolke *et al*. 2007). After cellularization has occurred additional oocyte growth is from import of yolk lipoproteins that are produced in the intestine (Greenstein 2005). We wanted to know if the change in the overall size of oocytes was driven primarily by increased cytoplasmic stream during moderate temperature stress. Thus, we examined the size of the last oocyte in the line (the one most recently cellularized) versus the first oocyte in the line (the one next to be ovulated) (Figure 5C). We found that in wild type worms there was a significant size difference between the first and last oocyte at 20°C, while at 26°C there was no significant difference in size between the first and last oocyte. There was a similar pattern in the sizes of the first and last oocytes in the *lin-54(n2231)* as seen in wild type. However, in *lin-35(n745)* mutants there was a similar difference in size between the first and last oocyte at both temperatures (Figure 5C). The last oocyte, but not the first oocyte, in *lin-35(n745)* mutants alone was significantly smaller than the same oocytes class in wild type at both temperatures (Figure 5C). These data suggest that the overall increased size in oocytes seen during moderate temperature stress is primarily driven by oocytes being bigger at the point that they are first cellularized.

## DISCUSSION

Germ cells that make oocytes have the combined need to provide an intact genome and the cytoplasmic components to the one cell embryo. Increasing germline apoptosis during stress can aid in both processes. Here we found that components of the DREAM complex, LIN-35 and the Muv B core proteins LIN-54 and LIN-37, are necessary for the full increase in apoptosis seen during temperature stress. Similarly, repression of CED-9 protein function is necessary for the full increase in apoptosis seen during temperature stress. Finally, induction of germline apoptosis during temperature stress is completely abolished when LIN-54 is absent and CED-9 constitutively active, suggesting these two pathways work together to regulate apoptosis during moderate temperature stress. In addition, we found that the rate of cytoplasmic streaming and the size of oocytes increases in wild type during moderate temperature stress. These findings expand the known role of LIN-35 and CED-9 and add the Muv B core as central regulators of stress induced germline apoptosis.

### Activation of apoptosis during stress may rely on a combinatorial regulation of CED-9/*ced-9*

The core apoptosis machinery is present and active in the germline to regulate apoptosis. Interestingly, there appears to be little role of regulation of CED-9 expression or function during physiological germline apoptosis in non-stressed/non-damaged germlines (Figure 6) (Gumienny *et al*. 1999, this work). However, as with other stressors, repression of CED-9 protein function appears to be an important step in allowing for activation of increased apoptosis during moderate temperature stress. Like in DREAM complex mutants, the constitutively active *ced-9(n1950)* mutant showed a dampened activation of apoptosis during moderate temperature stress. However, simultaneous loss of the DREAM complex Muv B core protein LIN-54 and expression of constitutively active CED-9 resulted in a total loss of activation of apoptosis during moderate temperature stress. Therefore, we are proposing a model were the combinatorial regulation of both *ced-9* mRNA expression and CED-9 protein function lead to germline apoptosis activation during stress (FIGURE 6). LIN-35 has been shown to repress *ced-9* expression in response to starvation stress and the DREAM complex has been shown to bind to *ced-9* operon in embryos (Gumienny *et al*. 1999; Groetsch *et al*. 2017). Therefore, we propose that the DREAM complex represses expression of *ced-9* leading to a lower level of CED-9 protein. Through another pathway, the remaining CED-9 protein is functionally deactivated, likely by EGL-1 binding (Gartner *et al*. 2000; Aballay and Ausubel 2001; Schumacher *et al*. 2005; Salinas *et al*. 2006), leading to the release of CED-4 and activation of the canonical apoptosis pathway. One question yet to be answered is why more cells are dying during moderate temperature stress. On one hand, more permissive conditions for apoptosis could be broadly present throughout the germline leading to a somewhat stochastic and random increase in apoptosis. This would look like general ramp up of physiological apoptosis (Gartner *et al*. 2008). Recent work suggests that differences in cytoplasmic flux into/outoff the compartment where nuclie lie in the distal germline may play a role in determining which nuclei are selected for physiological apoptosis (Chartier *et al*. 2021). The increase in cytoplasmic streaming we see during moderate temperature stress may result in/be a consequence of changes in cytoplasmic flux with more nuclei having cytoplasm move into the rachis, leading to more of these nuclei undergoing apoptosis through a method similar to that of physiological apoptosis. On the other hand, moderate temperature stress may lead to a specific type of damage, such as DNA damage or asynapsis, and then these conditions may lead to apoptosis of those cells that are more damaged. Further investigations would allow for distinguishing between these options.

**Figure 6:**
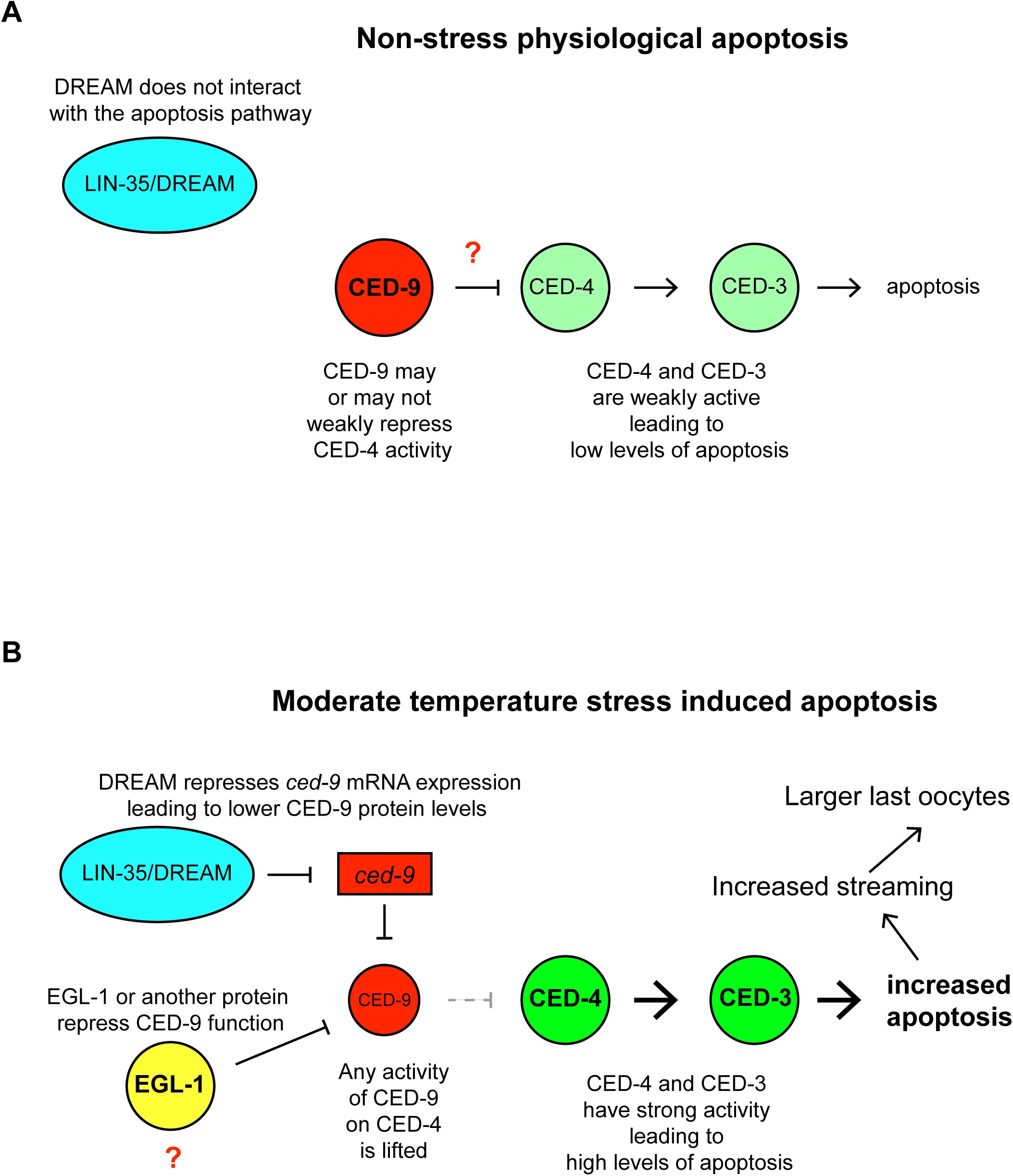
Model of apoptosis induction during moderate temperature stress. (A) Under non-stress conditions apoptosis levels are at physiological conditions. LIN-35 and the DREAM complex do not interact with apoptosis machinery. CED-9 does not play a large role in regulating the level of physiological apoptosis. (B) During moderate temperature stress LIN-35 and the DREAM complex likely repress the expression of *ced-9* leading to decreased levels of CED-9 proteins. Through EGL-1 or some other activity CED-9 binding of CED-4 is also relieved leading to CED-4 activation of the CED-3 caspase and increased levels of apoptosis. Higher rates of apoptosis lead to increased cytoplasmic streaming and larger oocytes right at the point of cellularization.

### The DREAM complex works with LIN-35 to activate apoptosis during stress

All of our data points towards a broad regulatory role of the DREAM complex, including the Muv B core and LIN-35, in the activation of germline apoptosis during stress. Previous work has shown that LIN-35 functions in activation of apoptosis in starvation stress and DNA damage situations (Schertel and Conradt 2007; Láscarez-Lagunas *et al*. 2014). We have added moderate temperature stress to this list. We also found that proteins in the Muv B core are equally responsible for activation of apoptosis during DNA damage and moderate temperature stress. While it seems likely that the DREAM complex functions to activate apoptosis through regulation of *ced-9* expression, what remains unclear is how these varied stressors are signaling to the DREAM complex and what entails activation of the DREAM complex in these situations. Some stressors such as DNA damage and asynapsis work through specific checkpoint proteins including CEP-1/p53 and BUB-3 respectively (Schumacher *et al*. 2001; Bhalla and Dernburg 2005). It is possible that multiple checkpoint proteins converge on the DREAM complex as the (one of the) central players in germline apoptosis activation. It is also unclear how stress leads to changes in DREAM complex function leading to changes in apoptosis. To date there have not been studies that looked at DNA binding pattern of the Muv B core of the DREAM complex in the germline. However, one study did look at areas of the genome associated with LIN-35 and the E2F/DP portions of the DREAM complex in non-stress germlines and found that the pattern of binding did not overlap extensively in the germline between them (Chi and Reinke 2006). These data suggest that at least under non-stress conditions LIN-35 and DP/EFL do not function together in the DREAM complex. Additionally, it has been shown that the E2F/DP and LIN-35 function on different levels of apoptosis regulation in the germ line, where LIN-35 regulates *ced-9* expression levels, while E2F/DP regulate *ced-4* expression levels (Schertel and Conradt 2007; Láscarez-Lagunas *et al*. 2014). Therefore, it may be possible that Muv B core functions with LIN-35 absent of E2F/DP in the germ line. The DREAM complex has been shown to be regulated by phosphorylation in mammalian systems leading to changes in DREAM complex formation and function (Litovchick *et al*. 2011; Guiley *et al*. 2015). The MAPK kinase pathway has been shown to be activated during increased apoptosis in response to other stressors including osmotic stress and heat shock (Salinas *et al*. 2006). Therefore, stress response pathways could converge on a kinase pathway, potentially MAPK kinase, to in some manner activate the entire DREAM complex (or just the Muv B core/LIN-35 portion) to repress *ced-9* expression contributing to activation of apoptosis.

### Chronic moderate temperature stress may allow animals to adapt and limit the cellular damage leading to apoptosis

In our experiments we saw there was a higher level of apoptosis induced in animals exposed to moderate temperature stress starting at the L4 stage when compared to animals where the moderate temperature stress started at the L1 stage. This suggests that whatever mechanism is disrupted in the germline leading to apoptosis may be able to adapt to chronic stress. The mechanism that seems most likely to be affect by temperature leading to increased apoptosis is formation of the synaptonemal complex (SC). Formation of the SC has been shown to fail starting at 26.5°C, just 0.5 degrees above our treatment temperature (Bilgir *et al*. 2013; Rog *et al*. 2017). Subtle changes in SC formation could lead to activation of the synapsis checkpoint, one of a number of checkpoints whose activation leads to apoptosis (Bhalla and Dernburg 2005). Similar to our data with apoptosis, previous data has suggested that longer times under moderate temperature stress allows the formation of the SC to begin to adapt to the temperature stress (Bilgir *et al*. 2013). However, this previous work only investigated different lengths of moderate temperature stress starting at the L4 stage or later. Therefore, this data cannot inform us on how moderate temperature stress may differently affect animals for whom their entire germline development is under moderate temperature stress (starting at L1) compared to animals that are under moderate temperature stress starting once the germline is fully formed (starting at L4). Further work will need to be done to test if chronic developmental temperature stress allows the formation of SC to fully adapt to the elevated temperature. It is also interesting to note that our findings on the level of induction of apoptosis are discordant with data on moderate temperature stress and brood size. In these experiments chronic exposure to moderate temperature stress starting at the embryo or L1 stage has a significantly strong effect on brood size compared to a shorter exposure starting at the L4 stage (Mikeworth and Cherian). Thus, there are likely additional effects of moderate temperature stress on germline development that limit fertility but do not directly lead to induction of apoptosis.

### Changes in streaming may contribute to oocyte size during stress

Previous research has shown that a decrease in apoptosis led to decreased cytoplasmic streaming and embryonic size (Wolke *et al*. 2007; Fausett *et al*. 2021). We saw the concomitant increase in streaming during moderate temperature stress when apoptosis is increased in wild type. The increase in streaming seen in wild type correlated with an increase in oocyte size in the last cellularized oocyte during moderate temperature stress, suggesting that the increase in streaming leads to a larger volume in oocytes. However, the first oocyte, the oocyte that is next to be ovulated, was not bigger during moderate temperature stress compared to non-stress conditions. Since oocyte growth after cellularization is primarily driven by import of yolk lipoprotein (Greenstein 2005), the lack of growth in the first oocyte could represent either a lower transport of yolk during moderate temperature stress or an overall constraint on oocyte growth due to the somatic gonad precluding oocytes from expanding past a specific size. In either case, oocytes are likely getting a larger percentage of their cytoplasmic volume from germline cytoplasmic streaming during moderate temperature stress than during non-stress conditions. The higher percentage of germline sourced cytoplasm in oocytes could include larger amounts of organelles, ribosomes, and germline expressed mRNAs and proteins that could positively or negatively affect embryos formed during moderate temperature stress. We did not see a clear pattern of changes in cytoplasmic streaming and oocyte growth in the DREAM complex mutants. Part of this may be due to the intermediate level of apoptosis seen in the mutants during temperature stress. However, it is well documented that these mutants have other pleiotropic effects on germline function (Goetsch *et al*. 2017; Mikeworth *et al*. 2023), which could mask any direct correlation between apoptosis level, cytoplasmic streaming rate, and oocyte size.

